# Cold Tolerance is Governed by Diverse Genetic Mechanisms Including Those Regulated by NB-LRR-type Receptor Proteins in Arabidopsis

**DOI:** 10.1101/2022.01.18.476799

**Authors:** Dipak K. Sahoo, Chinmay Hegde, Madan K. Bhattacharyya

## Abstract

Cold stress adversely affects the growth and development of plants and limits the geographical distribution of crop plants. Accumulation of spontaneous mutations shapes the adaptation of plant species to diverse climatic conditions. Genome-wide association study of the phenotypic variation gathered by a newly designed phenomic platform with that of the single nucleotide polymorphic (SNP) loci across the genomes of 417 Arabidopsis natural variants collected from various geographical regions revealed 33 candidate genes for cold tolerance. Investigation of at least two independent mutants for 29 of these genes identified 16 cold tolerance genes controlling diverse genetic mechanisms. This study identified five genes encoding novel leucine-rich repeat domain-containing proteins, including three nucleotide-binding site leucine-rich repeat (NBS-LRR) proteins. Among the 16 identified cold tolerance genes, *ADS2* and *ACD6* are the only two cold tolerance genes identified earlier. The comparatively little overlap between the genes identified in this genome-wide association study of natural variants with those discovered previously through forward and reverse genetic approaches suggests that cold tolerance is a complex physiological process governed by a large number of genetic mechanisms.

**Short Summary:** Cold stress adversely affects the growth and development of plants and limits the geographical distribution of crop plants. Genome-wide association study of the phenotypic variation of Arabidopsis natural variants with that of the single nucleotide polymorphic loci followed by T-DNA insertion mutant analyses of 29 candidate genes led to assigning cold tolerance function for the first time to 14 genes including three nucleotide-binding sites leucine repeat region (NB-LRR) protein genes. The comparatively little overlap between the genes identified in this study with those discovered previously suggests that cold tolerance is governed by a complex network of multiple genetic mechanisms.

## INTRODUCTION

Globally, 13.4 billion hectares or one-third of the total land area is potentially suitable for arable agriculture. Unfortunately, because of abiotic stresses, only approximately one-ninth of the potentially arable land is ideal for crop production (Bruinsma, 2003). Severe weather conditions such as extreme cold, and substantial and extended precipitation (Rosenzweig et al., 2002; Li et al., 2019b), hailstorms (Sánchez et al., 1996), and heatwaves and droughts (Ciais et al., 2005; van der Velde et al., 2010) limit agricultural productivity worldwide. Abiotic stresses affect the farming of existing crop species and act as a significant barrier for the introduced new crops. During acclimatization, a change in the expression levels of many genes allows the adaption of plant species to unique geographical regions. For example, investigation of Arabidopsis ecotypes collected from broad geographical regions has revealed genes essential for adaptation (Hancock et al., 2011; Fournier-Level et al., 2011).

Cold stress is a significant abiotic stress that adversely affects plants’ growth and development and restricts crop plants’ geographical distribution. Plants are classified as either chilling (0-15 °C) or freezing (< 0 °C) tolerant. These two classes are not mutually exclusive, as chilling tolerant plants in a temperate climate can induce their freezing resilience after exposure to chilling or non-freezing temperatures during cold acclimation (Lyons and Breidenbach, 1981). Cold acclimation in plants is linked to biochemical and physiological changes resulting from altered gene expression as well as bio-membrane lipid composition and accumulation of small molecules (Thomashow, 1998; Yamaguchi-Shinozaki and Shinozaki, 2006; Sanghera et al., 2011). Cold tolerance is a multifaceted trait linked to numerous cell compartments and metabolic pathways regulated by reprogrammed gene expression (Hannah et al., 2005). Plants from tropical and subtropical regions lack such cold acclimation machinery and are sensitive to chilling stress. The molecular basis of cold acclimation and acquired freezing tolerance has been investigated extensively in plants like Arabidopsis and winter cereals.

Arabidopsis is an ideal model plant for dissecting genetic pathways involved in combating environmental stresses. The 1,001 Arabidopsis Genomes Project initiative led to the resequencing of 1,135 natural inbred lines collected from the native Eurasian/North African range and the recently colonized North America (Alonso-Blanco et al., 2016). Genome-wide association studies (GWAS) of these natural variants adapted to three diverse ecological environments are expected to facilitate the identification of genetic mechanisms for adapting Arabidopsis to distinct climatic conditions. Earlier, GWAS in a limited number of natural variants of Arabidopsis revealed candidate genetic loci for adaptation (Hancock et al., 2011; Fournier-Level et al., 2011). GWAS of natural Arabidopsis variants can identify candidate genes for physiological functions. The function of such candidate genes can then be validated by studying knockout mutants for these genes. There are over 260,000 individual mutant lines in the Arabidopsis community, among which one can identify knockout, and knockdown mutants for most of the 29,454 predicted protein-coding Arabidopsis genes (Alonso et al., 2003; O’Malley and Ecker, 2010; O’Malley et al., 2015). Recently, digital photo-based objective phenotyping for this model plant has also been established for high-throughput phenomic analyses (Manacorda and Asurmendi, 2018; Vasseur et al., 2018).

In this study, we have (i) developed a high-throughput digital photo-based objective phenotyping method for Arabidopsis rosette leaves; (ii) collected responses of 417 resequenced diverse Arabidopsis natural variants to prolonged cold-stress using this phenotyping system; (iii) conducted GWAS to identify candidate cold tolerance genes; and (iv) validated the functions of individual candidate cold tolerance genes by studying at least two independent Arabidopsis mutants for each gene. We identified 33 candidate genes involved in cold tolerance. Investigation of T-DNA insertion mutants for most of these 33 genes revealed 16 cold tolerance genes. Surprisingly only two of these genes, *ADS2* encoding an acyl-lipid desaturase and *ACD6* encoding a novel ankyrin protein termed accelerated cell death 6, were previously identified as the cold tolerance genes (Chen and Thelen, 2013; Lu et al., 2009). We identified a strong candidate gene encoding a MYB transcription factor, AtMYB42, for cold tolerance. Five LRR domain-containing genes including three NB-LRR genes with no similarity to previously identified NB-LRR cold-stress-related genes were also identified. Our results suggest that cold tolerance is a complex physiological process governed by many genetic mechanisms.

## RESULTS

### A high-throughput digital image-based two-dimensional (2D) phenotyping system for rosette leaves

Seeds of 417 Arabidopsis (*Arabidopsis thaliana*) accessions (Alonso-Blanco et al., 2016; Table S1) originating from diverse ecological regions (Fig. S1a) were obtained from the Arabidopsis Biological Research Center (ABRC), Ohio State University, Columbus, OH, USA. To facilitate sowing of a similar number of seeds among replications, we compared two types of agaroses (standard agarose and low melting NuSieve agarose) at several concentrations (0.6%, 0.45%, 0.3%, 0.15%, and 0.1%) to determine the condition in which seeds will remain suspended for an extended period of time. The 0.1% (w/v) NuSieve Agarose was the best medium for resuspending seeds that were surface-sterilized with 95% ethanol. To break the seed dormancy, the seeds were stratified in the agarose medium at 4°C in the dark for five days (Fig. 1a). After stratification, approximately 15 seeds from each ecotype were sown in individual cells of Plug Tray-288 (Growers Supply, IA, USA) filled with soil (Sungro Horticulture Professional Growing Mix, Hummert International, MO, USA) (Fig. 1b).

**Figure 1.**
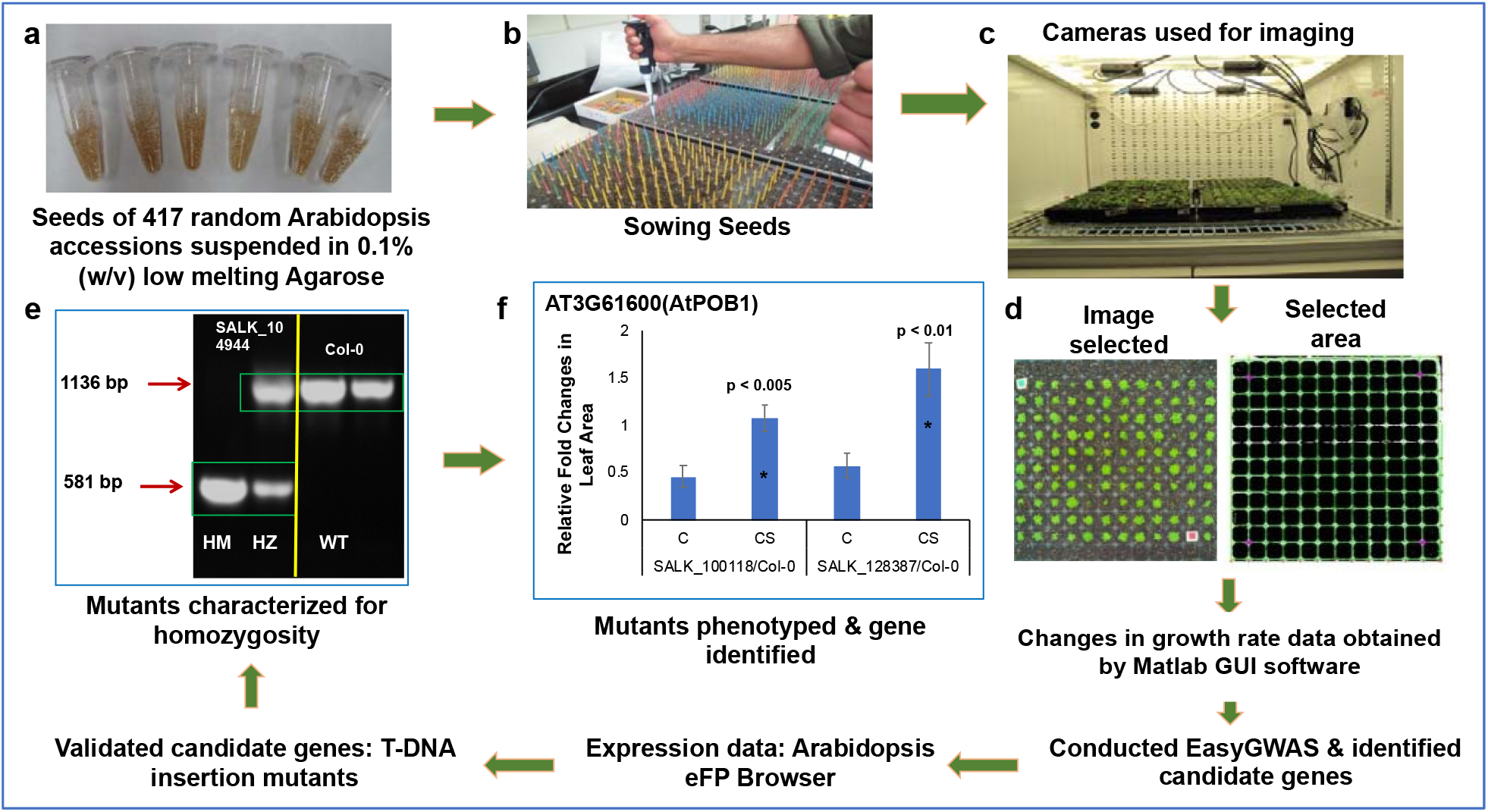
Schematic representation of the steps taken in identifying genes involved in cold stress using a high-throughput phenotyping platform. After suspending in low melting agarose **(a)**, seeds were stratified, and sown **(b),** and grown in the Arabidopsis growth chambers fitted with digital cameras **(c)**. Two-dimensional images of the rosette leaves of individual genotypes were captured and analyzed using Matlab GUI software (Method S2) **(d)**. The digital images were converted to pixel data for conducting easyGWAS to identify putative cold-stress-related genes. The candidate cold-stress-related genes were validated by studying knockout mutants. Homozygous knockout mutants were identified **(e)**. SALK_104944, knockout T-DNA insertion mutant for *AT1G68320*; HM, homozygous line; HZ, heterozygous line; WT (Col-0 ecotype), wild-type with no insertion. Knockout mutants were phenotyped to determine if any of the 33 putative cold-stress-related genes play a role in cold tolerance **(f)**.

The accessions were randomized within each of the three blocks in a randomized block design. The day of sowing the seeds was counted as “Day 0”. The plants were grown in AR-22L Arabidopsis chambers (Percival, IA, USA) (Fig. 1c). Four days after germination, seedlings were either thinned for a group of ten plants or transplanted to obtain one plant per cell. Plants were grown under 16 h day with light intensity 100 *μ*mol m^-2^ s^-1^, 22 °C temperature, 50% relative humidity (RH), and 8 h night with 18 °C temperature and 60% RH. The plants were watered once a week. On Day 7, in the “cold-stress” group, the temperature was reduced to 4 °C as the day and night temperature for 30 days, while plants in the “control group” continued to grow with no changes in growing conditions for seven additional days. Each tray contained the cold-tolerant accession PYL-6, the cold-sensitive accession Stepn-2, and the Columbia-0 (Col-0) ecotype with overlapping mutant lines to allow comparisons across an experimental group or between independent experiments to assure reproducibility. For each genotype, three experiments were conducted. In each experiment, 10 seedings were planted in 20 random cells in the “group of plants study.” Thus, each observation represented phenotypic datum collected from a group of ten plants grown in a cell; and for each genotype, data were collected from 60 cells (n = 60 from three experiments).

Two-dimensional (2D) images of the rosette leaves of a single or group of plants were captured by CropScore cameras (Computomics GmbH, Tübingen, Germany) during the day (Fig. 1d) and stored in the CropScore server (http://www.cropscore.com/en/home.html) for further analysis. A user-friendly software program Matlab GUI was written in Matlab to (i) capture and store a large dataset of high-resolution 2-dimensional (2D) digital images of aerial views of Arabidopsis rosette leaves in growth chambers over a period of time until leaves of the adjacent single or clusters of plants start to overlap; (ii) automatically crop, register and segment high-resolution images into sub-images corresponding to individual accessions based on a sequence of computer vision techniques (Method S1); (iii) extract important cues like rosette color and total rosette area from each sub-image, and store these cues in spreadsheet format for downstream statistical analysis (Method S2). Each of these stages is automatic, with an option included for manual intervention.

Our image processing workflow (Method S1-S2) executes the following six steps: (1) The Arabidopsis growth chambers are equipped with a network of point-and-shoot four cameras obtained from CropScore Inc (Computomics GmbH, Tübingen, Germany). Each Plug Tray-288 is divided into two halves, and each carrying 144 wells (12 x 12 wells) that are photographed by a single camera. Images are captured every 12 hours during the light period and transferred to a central server via an Ethernet connection (Fig. 1c). This step can be automated. (2) The images are cropped by detecting the tray boundaries if the camera is misoriented. This step can be automated. (3) For each image, the boundaries between adjacent cells in the grid are automatically detected using a two-step process (Fig. 1d): (i) each image is passed through a color filter tuned to the tray color, and (ii) edge boundaries, which are linear features in the filtered image, are estimated using a Hough transform. This step can be automated. (4) The 2D plane of the tray is estimated by calculating the geometric intersections of the cell boundaries. A perspective correction is applied to the original unfiltered image to register it to an orthographic view with respect to this plane. This step can be automated. (5) The registered image is now segmented into sub-images of different cells by simple cropping. This step can be automatic. (6) Cell images are passed through a second color filter tuned to the rosette leaves of healthy Arabidopsis. Rosette areas are then estimated by aggregating adjacent pixels, which have high responses to this color filter. This step can be made automatic. Steps 2 and 3 are the most challenging, and sometimes the automatic cropping and registration can fail if the tray boundaries are not detected correctly. If this is the case, the workflow signals the user to manually crop the image using a graphical user interface (GUI).

The utility of our software for two-dimensional (2D) Arabidopsis rosette leaf image analyses was evaluated through analysis of a set of random accessions (Fig. 2a) as follows. The images of 144 accessions were processed and analyzed by both FIJI (Schindelin et al., 2012; Abràmoff et al., 2004) and our Matlab software GUI (Method S1-S2). The 2D rosette leaf areas calculated by the two software for 144 accessions showed a statistically highly significant positive association (*r* = 0.99; *p* <0.01; Fig. 2a). Our Matlab GUI is a user-friendly software, and therefore we used this software to analyze the images. This approach’s key benefit is that it minimizes the amount of manual intervention and reduces data processing times by a factor of at least 20 compared to that for the existing commercially available phenotyping software solutions such as the FIJI software used in evaluating the performance of Matlab GUI (D.K. Sahoo, C. Hegde and M.K. Bhattacharyya, unpublished).

**Figure 2.**
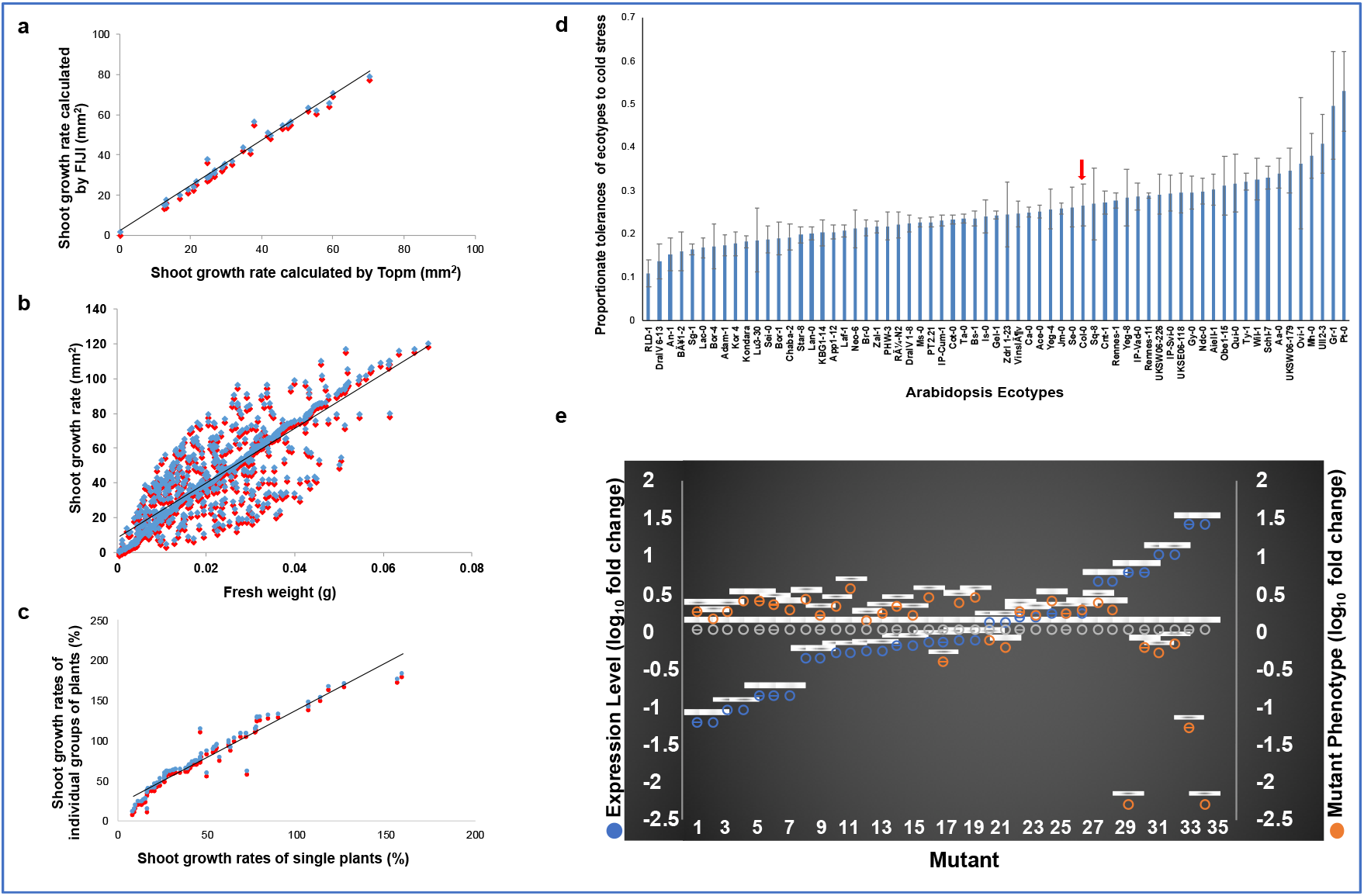
Phenotypic analyses of the cold tolerance trait in Arabidopsis. **(a)** Scatterplots demonstrating the relationships between the shoot surface area calculated by FIJI (blue) and Top.m (red) programs. Solid line displays statistically significant correlations (*r* = 0.99). **(b)** Relationships between the digital shoot surface area (sq. mm) calculated by Top.m (blue spot) and fresh weight of shoots. The scatterplot showed a positive association (*r* = 0.83) between the digital shoot surface area (sq. mm) calculated by Top.m (blue) and fresh weight (g) of shoots (red spot) among the 417 ecotypes. **(c)** Scatterplot demonstrating the positive association (*r* = 0.96) between the digital shoot surface area (%) from a group of plants (blue) with that of the corresponding single plant (red) of 76 randomly selected ecotypes. Growth Rate (%) in a-c was calculated as follows: [(Final Growth - Initial Growth)*100]/Initial Growth. **(d)** Tolerances of 417 Arabidopsis ecotypes to continuous cold stress. The growth rate of each ecotype (%) is calculated as = Growth on 30^th^ day of Cold Treatment X 100/Growth on 0^th^ Day of Cold Treatment. Proportionate tolerance of each ecotype is calculated as the growth rate of each ecotype X 100/ the summation of the growth rate of 417 ecotypes (detailed information is on Supplemental Fig. S2). The red arrow shows the proportionate growth of the ecotype Col-0. **(e)** Relationship of steady-state expression levels and mutant phenotypes of 16 cold tolerance genes. All but one knockout mutant of nine genes with reduced steady-state transcript levels under cold stress showed enhanced growth rates as compared to that in the wild-type ecotype Col-0 in response to prolonged low-temperature exposure (orange dots). Knockout mutants of four genes with enhanced steady-state transcript levels during cold stress showed reduced growth rates as compared to that in the wild-type ecotype Col-0 in response to prolonged cold stress. Grey dots showed log_10_ of 1 for transcript levels of genes or growth levels of mutants with no change at 4° as compared to that in wild-type Col-0. The data are from Table 1. Expression levels of individual genes (blue dots) at 24 h following exposure to cold stress (Figure S3) were used to plot the phenotypes of mutants identified for that gene.

We investigated if the 2D aerial images of Arabidopsis rosette leaves correctly predict the leaf growth of individual accessions. The 2D rosette leaf area of 417 accessions was determined using our software. The fresh weights (g) of rosette leaves of the same 417 accessions were also measured. The correlation coefficient between the 2D leaf area and fresh leaf weight (g) among the 417 accessions was *r* = 0.83 (*p*<0.01; Fig. 2b), suggesting the suitability of the phenotyping system for Arabidopsis at the seedling stage.

The growing of single Arabidopsis plants is labor-intensive. Therefore, we investigated if the 2D leaf area of ~15 plants can predict the 2D leaf area of a single plant. The association between the 2D image-based growth of single plants with that of groups of ~15 plants among 76 accessions was found to be highly significant (r = 0.96; *p*<0.01; Fig. 2c), suggesting that investigation of groups of plants instead of single plants should provide a reliable fresh weight estimate for single plants.

### Responses of Arabidopsis ecotypes to prolonged cold-stress

Using our 2D aerial rosette leaf phenotypic system, we investigated the responses of 417 Arabidopsis accessions, genomes of which have been resequenced (Alonso-Blanco et al., 2016), to prolonged cold-stress (4 °C). The 417 accessions include accessions studied previously for responses to non-freezing (Barah et al., 2013) and freezing cold stresses (Zhen and Ungerer, 2008; Xie et al., 2019). We collected the estimated 2D aerial rosette leaf areas of the selected 417 accessions under either 22° C (C) or prolonged cold stress at 4 °C (CS) in square mm (Table S2; Fig. S2). A 10-fold difference in the 2D aerial leaf area of the most cold-tolerant accession PYL-6 with that of the most cold-sensitive accession Stepn-2 was observed (Fig. 2d; Fig. S2). The broad-sense heritability (*h^2^*) for the aerial rosette leaf phenotype was 84%, suggesting that the phenotypes are reliable.

### A genome-wide association study (GWAS) revealed 33 putative cold tolerance genes

The 417 lines considered for this study were resequenced previously, and single nucleotide polymorphisms (SNPs) for the entire genome were predicted by comparing its genome sequence with the reference genome sequence of Columbia-0 (Col-0) ecotype (Grimm et al., 2017). We conducted a GWAS of the phenotypic variation of the 417 accessions for responses to prolonged cold stress with the SNPs distributed across the entire genome (Grimm et al., 2017). Pixel data of the natural variants were log-transformed to facilitate reliable parametric tests. The association of the phenotypic data with SNP data was tested using either a (i) linear regression (LR) or (ii) an <u>e</u>fficient <u>m</u>ixed-<u>m</u>odel <u>a</u>ssociation e<u>x</u>pedited (EMMAX) model (Grimm et al., 2017). An arbitrary cutoff *p* values of –log10 ≥ 4.5 identified 33 genes (Kaler and Purcell, 2019). The GWAS was conducted for each of the 17 independent experiments (CS; Cold Stress) and the mean data of the 17 experiments (Table S3). Quantile-quantile (QQ) plots (Fig. S1b) suggested that the data are normally distributed. The frequency distribution of the 417 accessions for proportionate cold tolerance also exhibited a normal distribution (Fig. S1c). Of the 33 genes, 15 were identified when EMMAX model was used, and seven when LR model was used. Eleven genes were detected in both LR and EMMAX models (Table S4).

### Expression patterns of the 33 candidate cold tolerance genes

A large number of genes are transcriptionally regulated in response to cold stress (Winter et al., 2007). Therefore, we investigated the 33 candidate cold tolerance genes for their expression patterns using the eFP database (Winter et al., 2007). To our surprise, 32 of the 33 genes are regulated at the transcriptional level to some extent in response to cold stress (Fig. S3; Table S5). This observation suggests that at least some of the 33 genes could be involved in adapting Arabidopsis to prolonged cold stress.

### Analyses of insertion mutants identified 16 genes involved in cold tolerance

We investigated 64 homozygous T-DNA insertion- and one transposon-induced mutants for the 33 candidate cold tolerance genes. We were able to identify two or more homozygous insertion mutants for 29 genes with one mutant each for the remaining four genes (Fig. 1e-f; Fig. S4; Table 1; Table S6-S7). At least two mutants for each of the 16 of 29 genes showed significant differences in rosette leaf growth from the wild-type Col-0 following prolonged exposure to 4°C for 30 days (Table 1). We termed these 16 genes as cold tolerance genes. We were able to observe mutant phenotypes in only one mutant for each of the nine genes (Table 2; Table S7). These nine genes may govern subtle cold tolerance phenotypes. It will require additional studies, including complementation analysis, to understand their function in cold tolerance. We consider these nine genes as strong candidate genes for cold tolerance. For the remaining four of the 29 genes, both mutants failed to show any mutant phenotypes (Table S7).

**Table 1.**
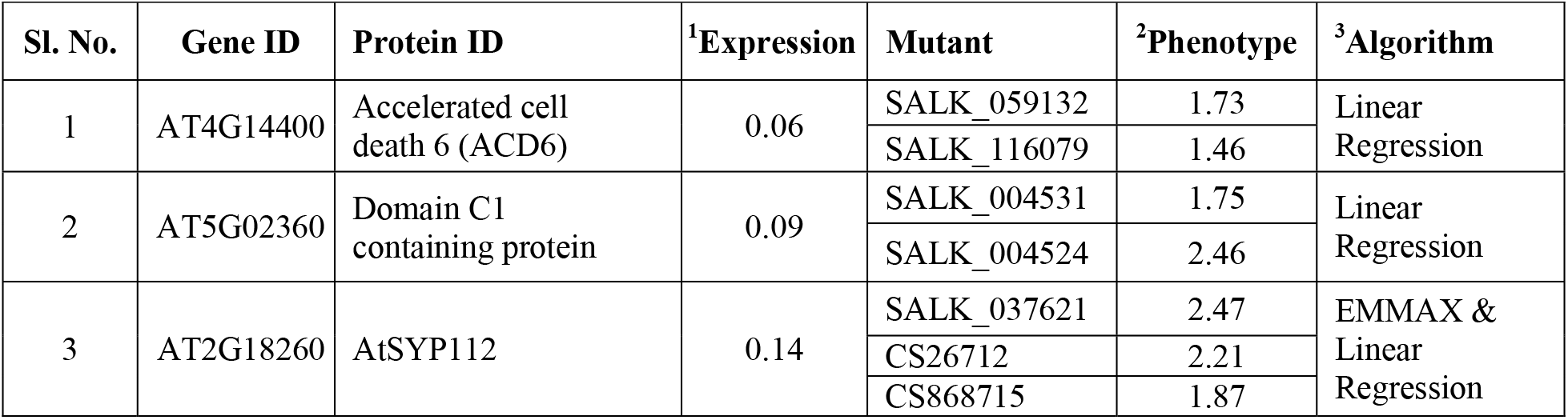
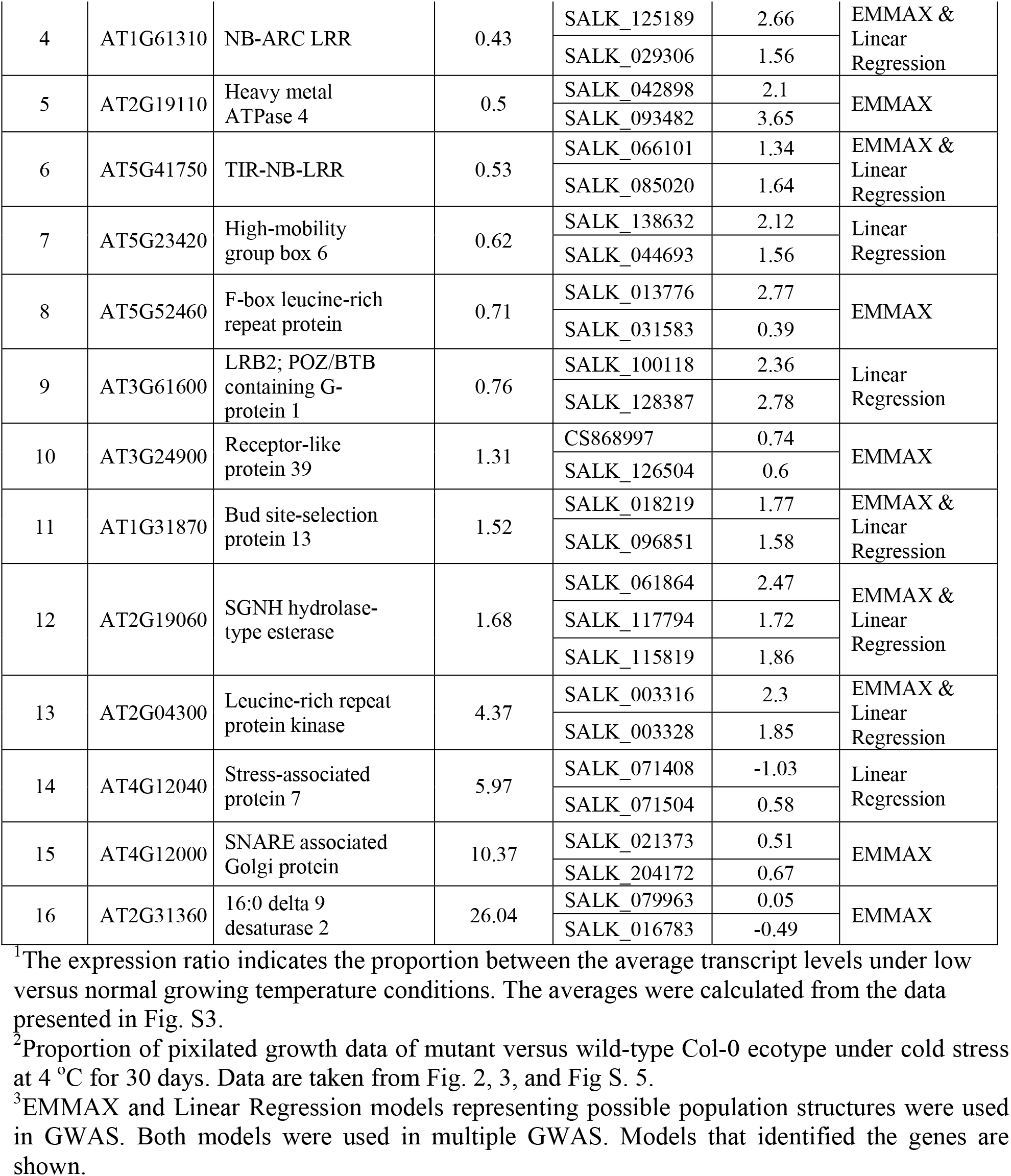
The 16 Arabidopsis genes involved in the expression of cold tolerance.

| Sl. No. | Gene ID | Protein ID | <sup>1</sup>Expression | Mutant | <sup>2</sup>Phenotype | <sup>3</sup>Algorithm |
| --- | --- | --- | --- | --- | --- | --- |
| 1 | AT4G14400 | Accelerated cell death 6 (ACD6) | 0.06 | SALK_059132 | 1.73 | Linear Regression |
|  |  |  |  | SALK_116079 | 1.46 |  |
| 2 | AT5G02360 | Domain C1 containing protein | 0.09 | SALK_004531 | 1.75 | Linear Regression |
|  |  |  |  | SALK_004524 | 2.46 |  |
| 3 | AT2G18260 | AtSYP112 | 0.14 | SALK_037621 | 2.47 | EMMAX & Linear Regression |
|  |  |  |  | CS26712 | 2.21 |  |
|  |  |  |  | CS868715 | 1.87 |  |
| 4 | AT1G61310 | NB-ARC LRR | 0.43 | SALK_125189 | 2.66 | EMMAX & Linear Regression |
|  |  |  |  | SALK_029306 | 1.56 |  |
| 5 | AT2G19110 | Heavy metal ATPase 4 | 0.5 | SALK_042898 | 2.1 | EMMAX |
|  |  |  |  | SALK_093482 | 3.65 |  |
| 6 | AT5G41750 | TIR-NB-LRR | 0.53 | SALK_066101 | 1.34 | EMMAX & Linear Regression |
|  |  |  |  | SALK_085020 | 1.64 |  |
| 7 | AT5G23420 | High-mobility group box 6 | 0.62 | SALK_138632 | 2.12 | Linear Regression |
|  |  |  |  | SALK_044693 | 1.56 |  |
| 8 | AT5G52460 | F-box leucine-rich repeat protein | 0.71 | SALK_013776 | 2.77 | EMMAX |
|  |  |  |  | SALK_031583 | 0.39 |  |
| 9 | AT3G61600 | LRB2; POZ/BTB containing G-protein 1 | 0.76 | SALK_100118 | 2.36 | Linear Regression |
|  |  |  |  | SALK_128387 | 2.78 |  |
| 10 | AT3G24900 | Receptor-like protein 39 | 1.31 | CS868997 | 0.74 | EMMAX |
|  |  |  |  | SALK_126504 | 0.6 |  |
| 11 | AT1G31870 | Bud site-selection protein 13 | 1.52 | SALK_018219 | 1.77 | EMMAX & Linear Regression |
|  |  |  |  | SALK_096851 | 1.58 |  |
| 12 | AT2G19060 | SGNH hydrolase-type esterase | 1.68 | SALK_061864 | 2.47 | EMMAX & Linear Regression |
|  |  |  |  | SALK_117794 | 1.72 |  |
|  |  |  |  | SALK_115819 | 1.86 |  |
| 13 | AT2G04300 | Leucine-rich repeat protein kinase | 4.37 | SALK_003316 | 2.3 | EMMAX & Linear Regression |
|  |  |  |  | SALK_003328 | 1.85 |  |
| 14 | AT4G12040 | Stress-associated protein 7 | 5.97 | SALK_071408 | -1.03 | Linear Regression |
|  |  |  |  | SALK_071504 | 0.58 |  |
| 15 | AT4G12000 | SNARE associated Golgi protein | 10.37 | SALK_021373 | 0.51 | EMMAX |
|  |  |  |  | SALK_204172 | 0.67 |  |
| 16 | AT2G31360 | 16:0 delta 9 desaturase 2 | 26.04 | SALK_079963 | 0.05 | EMMAX |
|  |  |  |  | SALK_016783 | -0.49 |  |

<sup>1</sup>The expression ratio indicates the proportion between the average transcript levels under low versus normal growing temperature conditions. The averages were calculated from the data presented in Fig. S3.
<sup>2</sup>Proportion of pixilated growth data of mutant versus wild-type Col-0 ecotype under cold stress at 4 °C for 30 days. Data are taken from Fig. 2, 3, and Fig S. 5.
<sup>3</sup>EMMAX and Linear Regression models representing possible population structures were used in GWAS. Both models were used in multiple GWAS. Models that identified the genes are shown.

**Table 2.**
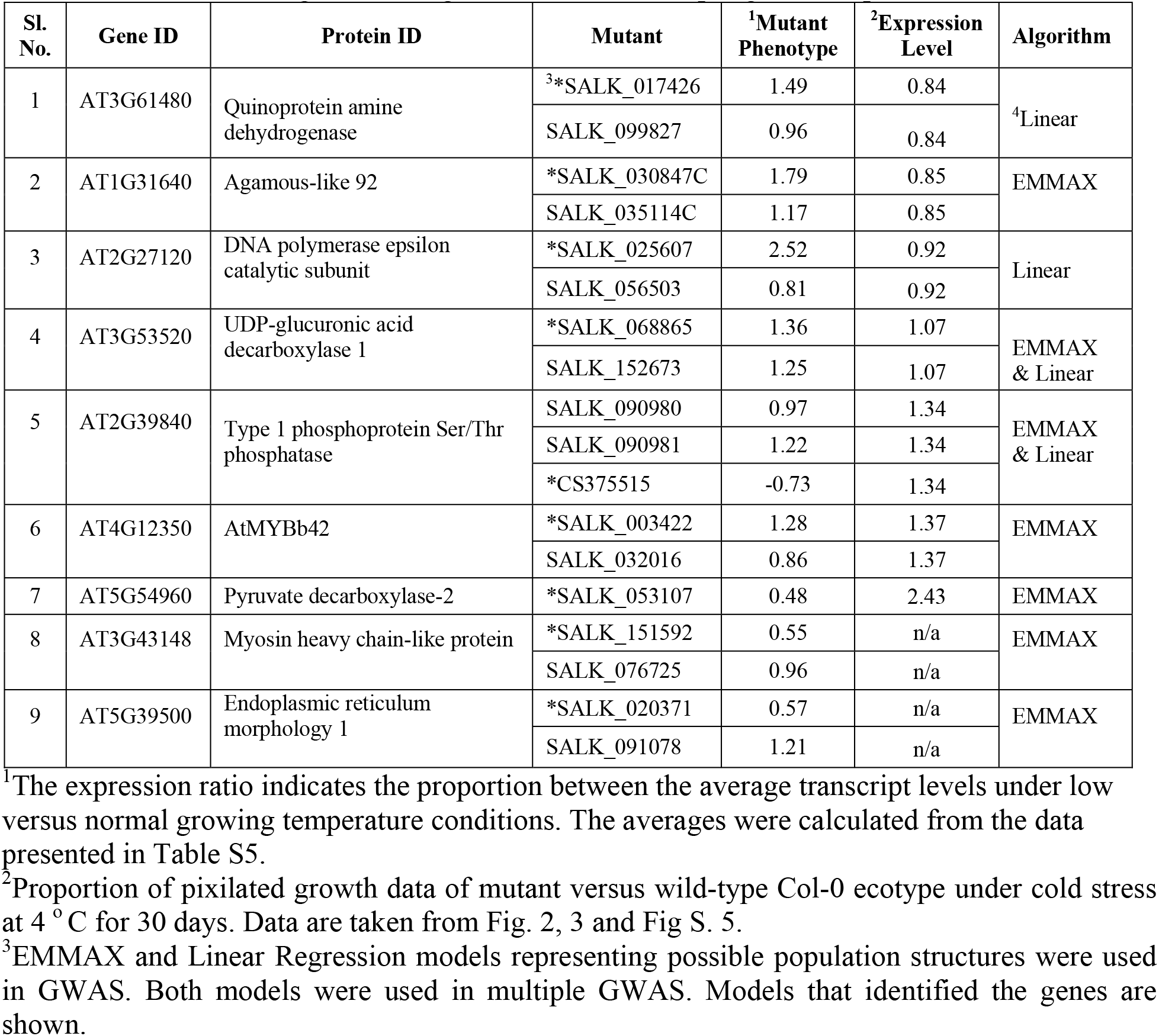
The nine strong candidate genes involved in adapting Arabidopsis to cold stress.

We investigated if there was any relationship between levels of transcripts and responses of the knockout mutants of the identified 16 genes to prolonged cold stress. We hypothesized that knockout mutants of the genes with reduced transcript levels under cold stress would have enhanced cold tolerance, while mutants of the genes that are induced during cold stress would show enhanced sensitivity to cold stress. Mean transcript levels for each of the 16 genes were calculated from six data points (Table 1; Fig. S3). Each of the 16 genes showed significant up- or down-regulation for at least one of the six data points (Winter et al., 2007; (Fig. S3). We did observe a clear relationship for thirteen of the 16 genes as expected; knockout mutants for nine genes with reduced transcript levels under cold stress showed enhanced cold tolerance while mutants of four genes with increased transcript levels under cold stress exhibited increased cold sensitivity (Fig. 2e; Table 1).

### Functions of the identified 16 cold tolerance genes

Interestingly only two of the 16 genes identified have previously been reported as cold tolerance genes: *AT4G14400*, which conditions accelerated cell death 6 (ACD6), and *At2g31360*, which encodes an acyl-lipid desaturase 2 (ADS2). *ACD6* (*At2g31360*) is downregulated by cold stress, and its loss of function knockout mutants showed enhanced cold tolerance to cold stress (Fig. 3; Table 1; Fig. S3a). This gene has been shown to be involved in chilling and freezing tolerance (Chen and Thelen, 2013). A gain of function *acd6* mutant shows an increased accumulation of salicylic acid level and exhibits freezing sensitivity (Lu et al., 2009; Miura and Ohta, 2010; Chen and Thelen, 2013) along with enhanced resistance against both bacterial and oomycete pathogens (Todesco et al., 2010).

**Figure 3.**
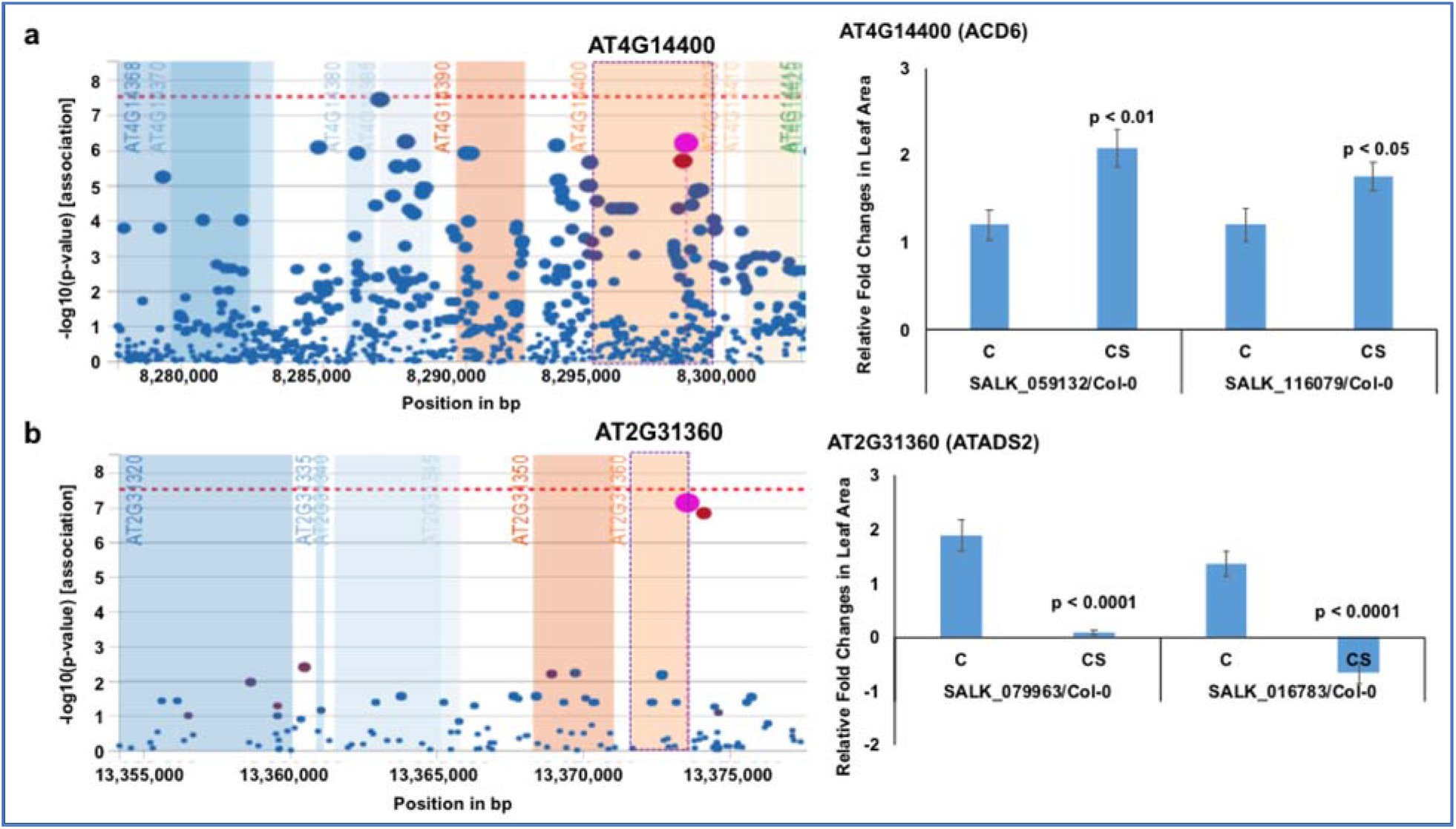
Previously identified two proteins that contribute either negatively or positively to cold tolerance. (a) *AT4G14400* encoding Accelerated cell death 6 (ACD6) protein negative regulates cold tolerance. (b) *AT2G31360* encoding acyl-lipid desaturase 2 (ADS2) positive regulates cold tolerance. On the left, output plot of *p*-values (-log base 10) in a 5-kb window for association of SNPs with phenotypic variation, obtained from easyGWAS is presented. On the right, rosette leaf growth rates of mutants with respect to Col-0 are presented. The relative rosette leaf growth rate in the mutant compared to wild-type Col-0 is significantly different in control (C) and cold stress (CS) (*p* < Bonferroni adjusted α) conditions. Knockout mutation in the *ADS2* gene resulted in yellowing and death of plants resulting in a negative growth rate in one T-DNA insertion mutant at 4°C and as compared to that at 22°C. C, Comparative growth rate of the mutant relative to wild-type Col-0 in control condition; CS, Comparative growth rate of the mutant relative to wild-type Col-0 in cold stress.

*ADS2* (*At2g31360*) is upregulated by cold stress, and the *ads2* mutant was shown to display a dwarf and sterile phenotype in response to cold stress at 6 °C and to show increased sensitivity to freezing temperature (Chen and Thelen, 2013). Here we also observed that the mutants of the cold-induced *ADS2* gene showed increased sensitivity to cold stress (Fig. 3b; Table 1; Fig. S3b). In *ads2* mutant plants, the membrane lipid composition is altered as compared to the wild-type. Reduced levels of 16:1, 16:2, 16:3, and 18:3 lipids and higher levels of 16:0 and 18:0 fatty acids were detected in the *ads2* mutant compared to the wild-type. The paralogous *acyl-lipid desaturase 1* and *2* (*ADS1* and *ADS2*) genes are induced in response to cold stress to facilitate cold acclimation and chilling/freezing tolerance, respectively (Chen and Thelen, 2016, 2013). *ADS1* encodes a soluble Δ9-desaturase that is found primarily in the chloroplast and catalyzes the desaturation of stearic acid (18:1) of monogalactosyl diacylglycerol (MGDG) (Barrero-Sicilia et al., 2017; Berestovoy et al., 2020); while, *ADS2* encodes a 16:0 desaturase for synthesis of MGDG and phosphatidylglycerol (Chen and Thelen, 2013). Both genes affect chloroplast membrane desaturation and have been shown to be essential for the cold adaptation response in Arabidopsis (Barrero-Sicilia et al., 2017). Reidentification of *ACD6* and *ADS2* cold tolerance genes validate our approach of identifying cold tolerance genes using a novel phenotyping system for Arabidopsis seedlings.

Among the 14 cold tolerance genes identified in this study, five contain LRR domains with unknown functions (Fig. 4; Table 1). *AT1G61310* encodes an LRR and NB-ARC domains-containing disease resistance-like protein, while *AT5G41750* encodes a TIR-NBS-LRR-type disease resistance-like protein. *AT5G52460* encodes an F-box leucine-rich repeat protein, annotated as embryo sac development arrest 41 (EDA41). The transcript levels of all three genes were suppressed by cold stress (Fig. S3c-e). All knockout mutants except one for these three genes showed enhanced cold tolerance as compared to the wild-type Col-0 ecotype; one T-DNA insertion mutant, SALK_031583, for the *AT5G52460* gene showed cold sensitivity (Fig. 4a-c; Table 1). In the SALK_031583 mutant, T-DNA was inserted in the promoter region, which might have enhanced the transcription of the gene leading to increased cold sensitivity. *AT2G04300* encodes an LRR protein kinase, and *AT3G24900* encodes the receptor-like protein 39 (AtRLP39) containing an LRR domain. The transcript levels of these two genes are induced during cold stress (Fig. 4d-e; Table 1; Fig. S3f-g). Surprisingly, although the *AT2G04300* gene is highly induced, the knockout mutants showed increased growth instead of reduced growth as compared to the control wild-type Col-0 ecotype under prolonged cold stress (Fig. 4d; Table 1; Fig. S3f). The knockout mutants of the cold stress-induced gene *AT3G24900* exhibited reduced growth as compared to that in the wild-type Col-0 ecotype under prolonged cold stress (Fig. 4e; Table 1; Fig. S3g).

**Figure 4.**
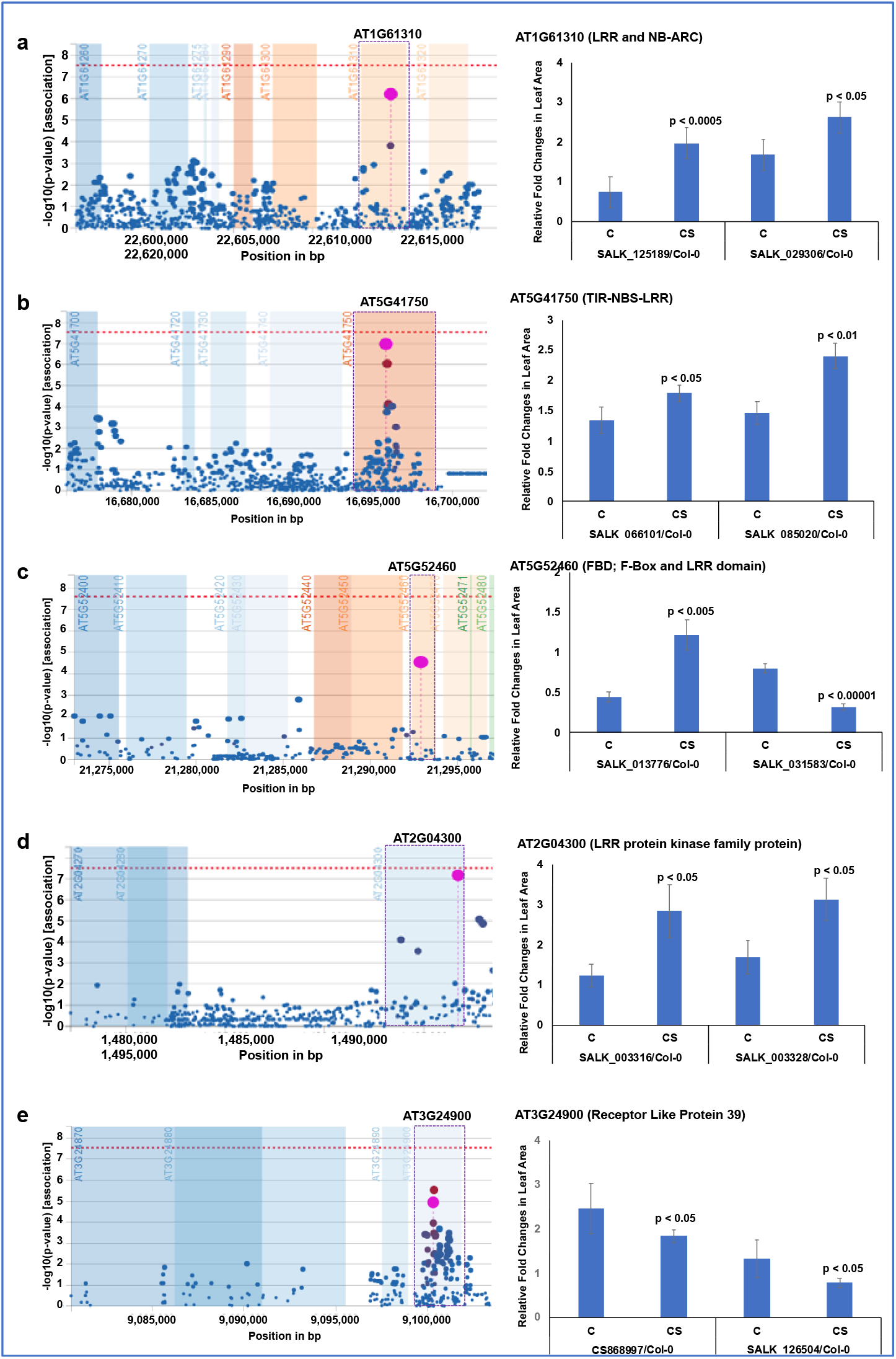
Five leucine-rich repeat domain-containing proteins contribute towards cold tolerance. **(a–e)** Each of the five proteins contains LRR domain. On the left, output plot of *p*-values (-log base 10) in a 5-kb window for association of SNPs with phenotypic variation, obtained from easyGWAS is presented. On the right, rosette leaf growth rates of mutants with respect to Col-0 are presented. The relative rosette leaf growth rate in the mutant compared to wild-type Col-0 is significantly different in control (C) and cold stress (CS) (*p* < Bonferroni adjusted α) conditions. C, Comparative growth rate of the mutant relative to wild-type Col-0 in control condition; CS, Comparative growth rate of the mutant relative to wild-type Col-0 in cold stress.

In addition to assigning cold tolerance functions to five novel LRR domain contacting proteins, we assigned cold tolerance function to nine additional genes. They are: (i) *AT1G31870* encodes the bud site-selection protein 13 (AtBUD13), which is involved in pre-mRNA splicing and embryo development (Xiong et al., 2019); (ii) *At5g02360* encoding DC1 domain-containing protein, (iii) syntaxin of plants 112, (iii) heavy metal ATPase 4, (iv) high-mobility group box (HMGB) 6, (v) LRB2; POZ/BTB containing G-protein 1, (vi) Bud site-selection protein 13, embryo sac development arrest 41 protein, (vii) SGNH hydrolase-type esterase, (viii) stress-associated protein 7, and (ix) SNARE associated Golgi protein (Table 1).

*AT1G31870* encodes the bud site-selection protein 13 (AtBUD13) (Table 1; Fig. S3h; Fig. S5a), which is involved in pre-mRNA splicing of 52 genes, of which 22 are involved in nucleic acid metabolism and embryo development (Xiong et al., 2019). Cold stress alters the expression and splicing of serine/arginine-rich (SR) genes that encode splicing factor proteins required for constitutive and alternative splicing (Leviatan et al., 2013; Palusa et al., 2007). *AtBUD13* could be a regulatory gene that could control cold-stress-related genes for cold adaptation through splicing.

Transcription of *AT2G18260* gene encoding syntaxin protein ATSYP112 also suppressed by cold stress loss of function mutants are cold tolerant; and therefore, we consider that this protein negatively contributes towards cold tolerance in Arabidopsis (Table 1; Fig. S3i; Fig. S5b).

*AT2G19060* encodes an SGNH hydrolase-type esterase (Table 1; Fig. S3j; Fig. S5c). The GDSL esterases/lipases are mainly involved in regulating plant development, morphogenesis, synthesis of secondary metabolites, and defense response (Kwon et al., 2009; Hong et al., 2008; Chepyshko et al., 2012). The GDSL family is further classified as SGNH hydrolase because of the presence of the strictly conserved residues Ser-Gly-Asn-His in the conserved blocks I, II, III, and V (Chepyshko et al., 2012). The role of GDSL family esterase in cold adaptation was reported in *Photobacterium sp*. strain J15 (Shakiba et al., 2016). Here, we reported a SGNH hydrolase-type esterase as a negative regulator during the cold stress response.

*AT2G19110* encodes the Arabidopsis heavy metal ATPase 4 (AtHMA4) with similarity to Zn ATPase (Meyer et al., 2016; Lekeux et al., 2018). Transcription of this gene is downregulated by cold stress and knockout mutants show enhanced cold tolerance suggesting a negative role of *AtHMA4* in cold tolerance (Table 1; Fig. S3k; Fig. S5d). *ATHMA4* is involved in the hyperaccumulation of Zn/Cd (Lekeux et al., 2019). Presumably, metal ion accumulation may be detrimental during cold stress. A study conducted in the halophyte four-wing saltbush (*Atriplex canescens*) revealed a heavy metal-associated protein, AcHMA1, whose expression was strongly downregulated under NaCl stress and cold stress (Sun et al., 2014). In Arabidopsis, such a mechanism might be mediated by *ATHMA4*, down-regulated under cold stress.

*AT3G61600* encodes the LIGHT-RESPONSE BTB2 (LRB2) protein, which together with LRB1, negatively regulates the photomorphogenesis (Christians et al., 2012). This gene is involved in protein ubiquitylation through interacting with CULLIN 3 proteins (Kim et al. 2016; Ni et al., 2014; Hu et al., 2014; Christians et al., 2012; Gingerich et al., 2005). In winter-annual accessions of *A. thaliana*, cold stress exposure or vernalization is needed to commence flowering via FRIGIDA (FRI). FRI acts as a scaffold protein to recruit numerous chromatin modifiers that epigenetically modify flowering genes and regulate blooming via proteasome-mediated degradation of FRI. During vernalization, FRI directly interacts with the BTB proteins LRB1/2, as well as the CULLIN3A (CUL3A) ubiquitin-E3 ligase *in vitro* and *in vivo* leading to proteasomal degradation of FRI (Christians et al., 2012; Hu et al., 2014). Long-term cold stress accelerates the degradation of FRI and blooming by decreasing FLC transcription, a mechanism dependent on CUL3A and associated with long non-coding RNA and chromatin remodeling (Hu et al., 2014). Here we have shown that the transcription of *LRB2* is suppressed by cold stress and knockout mutants showed enhanced cold tolerance suggesting its negative function for cold tolerance (Table 1; Fig. S3l; Fig. S5e).

The *AT4G12000* gene encodes a member of the soluble N-ethylmaleimide-sensitive factor adaptor receptor (SNARE)-associated Golgi protein family (Xu et al., 2019). Arabidopsis, 53 genes have been annotated to encode SNARE molecules (Sanderfoot et al., 2000). SNARE proteins in plants are involved in various physiological processes, including abscisic acid-related signaling and osmotic stress tolerance (Bassham and Blatt, 2008; Uemura et al., 2004) and soluble N-ethylmaleimide-sensitive factor adaptor protein (SNAP) genes are induced at low temperature (Bao et al., 2008). The expression of *OsSNAP32* was dramatically increased in rice seedlings treated with H_2_O_2_, PEG6000, and low temperature or after inoculation with the rice blast pathogen *Megnaporthe oryzae*. A gene family encoding SNAP25-type proteins is induced in rice following exposure to biotic and abiotic stresses (Bao et al., 2008). *AT4G12000* encoding a SNARE-associated Golgi protein identified in the present study is also induced under cold stress, and loss of function mutants for this gene displayed cold-sensitive phenotypes suggesting a positive function for this gene in cold tolerance (Table 1; Fig. S3m; Fig. S5f).

*AT4G12040* encodes an A20/AN1-like zinc finger family protein, stress-associated protein 7 (ATSAP7) (Vij and Tyagi, 2006). A20/ANI zinc-finger domain-containing SAPs are involved in abiotic stress (Mukhopadhyay et al., 2004). Another SAP, AtSAP10, is involved in various abiotic stresses such as heavy metals and metalloids, high and low temperatures, and treatment with ABA (Dixit and Dhankher, 2011). *AtSAP12* is induced following cold treatment (Ströher et al. 2009). Expression of both *OsiSAP1* and *OsiSAP8* is induced in rice in response to a variety of environmental stresses, including cold, drought, heavy metals, wounding, and submergence (Mukhopadhyay et al., 2004; Ae and Gupta, 2008) and overexpression of *OsiSAP8* provides rice with strong tolerance to cold, salt, and drought (Mukhopadhyay et al., 2004; Ae and Gupta, 2008). Overexpression of *ZFP177*, a rice zinc-finger *A20/AN1* gene, in tobacco resulted in enhanced tolerance to both high and low temperature (Huang et al., 2008). Similarly, overexpression of AlSAP, a stress-associated protein from a halophyte grass *Aeluropus littoralis*, in tobacco provides increased tolerance to cold, heat, salt, and drought stresses (Saad et al. 2010). Expression of *ATSAP7* (*AT4G12040*) is induced by cold stress and both T-DNA insertion knockout mutants identified for this gene exhibited reduced growth rate under prolonged cold stress as compared to that in the wild-type Col-0 ecotype, a positive cold tolerance function as observed for other Arabidopsis and rice SAP proteins containing A20/AN1-like zinc finger domains (Table 1; Fig.3n; Fig. S5g).

*At5g02360* encodes a cysteine/histidine-rich divergent C1 (DC1) domain-containing novel protein (Allen et al., 2004; Fig. S5h). The transcription of this gene is suppressed in response to cold stress (Fig. S3o). A cysteine/histidine-rich DC1 protein has been shown to have a positive regulatory function in cell death and plant defenses in pepper (Hwang et al., 2014). Here we have shown that knockout mutants for *At5g02360* are highly tolerant to cold stress compared to the wild-type control, suggesting a possible negative function for this gene (Table 1; Fig.3o; Fig. S5h).

*AT5G23420* (Table 1; Fig. S3p; Fig. S5i) encodes a high-mobility group box 6 (HMGB6) protein (Pedersen and Grasser, 2010; Kwak et al., 2007; Grasser et al., 2004). HMGB nuclear proteins are involved in various cellular processes including replication, transcription, and nucleosome assembly. The Arabidopsis genome contains eight genes encoding HMGB proteins (Grasser et al., 2004; Kwak et al., 2007). Cold treatment increases the expression of *HMGB2, HMGB3*, and *HMGB4*, whereas the transcript levels of *HMGB1* and *HMGB5* are not noticeably affected by cold stress (Kwak et al., 2007). The expression of *AT5G23420* is suppressed by cold stress, and both T-DNA insertion knockout mutants for *AT5G23420* exhibited enhanced cold tolerance (Table 1) suggesting a negative function for this protein in cold tolerance.

Several genes have been identified as cold stress regulatory and responsive genes through forward genetic screening. Among the 16 genes we have identified as cold tolerance genes, only *ACD6* and *ADS2* were previously shown to be freezing responsive genes (Chen and Thelen, 2013). *AT4G12350* gene encoding ATMYB42 (Table 2), with a subtle cold stress-related phenotype, is a homologue of the ATMYB14/15 transcription factors that have been demonstrated to negatively regulate at least some cold stress response genes (Agarwal et al., 2006; Chen et al., 2013; Miura et al., 2007). ATMYB42 has been shown to be a regulator of phenylalanine and lignin biosynthesis (Geng et al., 2020).

Blast2GO analysis was conducted to understand the functional annotation of the 16 cold tolerance genes (Conesa et al., 2005; Götz et a. 2008) (Fig. S6; Table S8-S10). The 16 genes were grouped into 58 classes based on their biological processes, to 17 classes according to their molecular functions, and to 13 classes as cellular components or based on their subcellular locations suggesting that most, if not all, of the 16 cold tolerance genes encode multiple functions (Fig. S6; Table S8-S10). Collectively, alterations in genes associated with defense response (*AT1G61310, AT4G14400*), metabolic processes including nucleic acid metabolic process (AT1G31870), ubiquitin-dependent protein catabolic process (*AT4G12040*), protein ubiquitination (*AT3G61600, AT4G14400*), lipid metabolic process (*AT2G19060, AT2G31360*), transport (*AT2G18260*) including ion transmembrane transport (*AT2G19110*), organic cyclic compound metabolic process (*AT1G31870*) suggest the complexity of genetic mechanisms involved in cold tolerance. Blast2GO (Conesa et al., 2005) functional analysis also identified two cold-responsive genes (*AT2G18260* and *AT4G14400*) (Fig. S5a; Table. S8).

The Kyoto Encyclopedia of Genes and Genomes (KEGG) (Kanehisa et al., 2017; Götz et a. 2008) pathway analyses of 25 genes including nine genes with subtle cold tolerance phenotypes revealed that *AT2G18260* is involved in the pantothenate and CoA biosynthesis pathway (EC:2.7.7.3 in Fig. S7), and *AT2G31360* in the biosynthesis of unsaturated fatty acids (EC:1.14.19.1 in Fig. S7) (Table 1). The metabolome of Arabidopsis under temperature stress showed an increase in a small group of amine-containing metabolites (β-alanine and putrescine) (Kaplan et al., 2004), and plants capable of cold acclimation accumulate polyunsaturates during cold stress. KEGG pathways analysis revealed that *AT2G27120* (Table 2) is involved in DNA replication, base/nucleotide excision repair, and purine metabolism pathways (EC:2.7.7.7 in Fig. S7), *AT3G53520* (Table 2) is involved in amino sugar and nucleotide sugar metabolism pathways (EC:4.1.1.35 in Fig. S7), and *AT5G54960* (Table 2) involved in glycolysis/gluconeogenesis (EC:4.1.1.1 in Fig. S7). The connection between plant DNA damage response and responses to biotic and abiotic stresses has been reported (Nisa et al., 2019). Duplication of genes has been observed specifically for those involved in reproduction, DNA damage repair, and cold tolerance in the high-altitude plant, *Eutrema heterophyllum* (Guo et al., 2018).

### MapMan analysis of cold-responsive genes

The efficacy of the full-genome sequences for important crop species has been advanced by the development of detailed ontologies by programs such as MapMan (Usadel et al., 2009; Schwacke et al., 2019), which has assigned enzymes to over 1,200 groups covering almost all central metabolic pathways. Mapman facilitates analyses of large transcriptomic and proteomic datasets (Usadel et al., 2009). Though it was developed initially for analyses of Arabidopsis datasets (Usadel et al., 2009), MapMan ontology has been extended to several other species (Ling et al., 2013) including soybean (Nanjo et al., 2011), cotton (Christianson et al., 2010), maize (Usadel et al., 2009), potato (Kondrák et al., 2011), and tomato (Barsan et al., 2010). MapMan maps transcript profiling data onto pathways and genetic maps and generates response overlays that simplify the identification of shared features globally and on a gene-to-gene basis (Usadel et al., 2009; Pitzschke and Hirt, 2010). We conducted MapMan analyses of the identified 16 cold tolerance genes (Table 1) and nine strong candidate genes (Table 2) using their transcript profiles (Figs. S3 & Table S5) and mapped 23 of the 25 genes to metabolism, biotic stress, cellular response, proteasome, autophagy and cellular function categories (Fig. S8).

## DISCUSSION

Genetic pathways regulating the expression of cold stress-responsive genes have been identified through either forward (Provart et al., 2016) or reverse genetics (Provart et al., 2016; Chinnusamy et al., 2010; Østergaard and Yanofsky, 2004; Alonso and Ecker, 2006). In *Arabidopsis*, changes in gene expression in response to cold stress are regulated by the C-REPEAT BINDING FACTOR (CBF)-mediated cold signaling pathway (Chinnusamy et al., 2010; Jeon and Kim, 2013). Cold stress elevates Ca^2+^ levels transiently and activates protein kinases, including MAP kinases, for cold acclimation (Lissarre et al., 2010). In transgenic Arabidopsis, overexpression of CBF/DREB proteins led to desiccation and cold tolerance through ectopic expression of RD/COR genes (Jaglo-Ottosen et al., 1998; Kasuga et al., 1999). The transcription factors CBF (C-repeat-binding factor)/DREB1 (dehydration responsive element binding1) and ICE1 (inducer of CBF expression 1) have essential roles in regulating the expression of cold-responsive (COR) genes (Chinnusamy et al., 2010; Lissarre et al., 2010; Liu et al., 1998). The CBFs/DREBs induce several hundred genes by binding to their CRT/DRE elements (Vogel et al., 2005). REIL2 deficiency delays CBF/ DREB regulon activation and reduces CBF/ DREB protein accumulation in response to cold stress (Yu et al., 2020). Overexpression of a ribosomal biogenesis factor encoded by *STCH4/REIL2* enhances chilling and freezing tolerance in Arabidopsis (Yu et al., 2020). STCH4 presumably induces alterations in the ribosomal composition and functions at low temperatures to facilitate the translation of proteins required for plant development and survival under cold stress (Yu et al., 2020). Likewise, overexpression of genes encoding ice recrystallization inhibition (IRI) proteins LpIRI-a or LpIRI-b in Arabidopsis exhibited improved cell membrane stability in freezing and improved frost tolerance (Zhang et al., 2010). Open stomata 1 (OST1) protein kinase also plays a central role in regulating freezing tolerance in Arabidopsis and its activity is regulated by a plasma membrane-localized clade-E growth-regulating 2 (EGR2) phosphatase (Ding et al., 2019).

In this study, we investigated Arabidopsis natural variants collected from a broad geographical region to identify possible additional genetic mechanisms for of cold tolerance. A diverse collection of natural variants is a useful resource for identifying genetic mechanisms involved in various biological processes. Arabidopsis is an ideal model plant distributed across 15.11 to 62.63 latitudes and −123.21 to 136.31 longitudes that include a diverse ecological range, including its ancestral Iberian Peninsula habitat to northern latitudes with an unknown glacial refugium (Alonso-Blanco et al., 2016). The 1,135 natural Arabidopsis variants collected from the diverse ecological ranges have been resequenced to facilitate identifying candidate genes for various traits through GWAS (Alonso-Blanco et al., 2016). The model plant Arabidopsis is particularly suitable for this study because of the availability of a large collection of mutants to validate the candidate genes identified in GWAS (O’Malley et al., 2015).

The genetic basis of cold tolerance in numerous crops has been investigated using genome-wide association mapping (GWAS). For example, a GWAS for cold tolerance at the seedling stage among rice landraces discovered a total of 26 SNPs that were significantly associated with cold tolerance (Song et al., 2018). Similarly, GWAS and differentially expressed gene (DEG) analysis among germinating maize seeds revealed 30 SNPs and two DEGs that were associated with cold tolerance (Zhang et al., 2020). A GWAS among 200 cotton accessions collected from diverse geographical locations revealed an alcohol dehydrogenase gene (*GhSAD1*) associated with cold tolerance (Ge et al., 2021). In canola, GWAS led to identification of 25 candidate genes that were previously shown to be associated with freezing tolerance, photosynthesis, or cold responsiveness in canola or Arabidopsis (Chao et al., 2021).

We developed a high-throughput phenotyping platform and determined the responses of seedlings of 417 of these ecotypes to cold stress at 4 °C for 30 days. The 417 ecotypes showed a 10-fold difference in growth rate between the most cold-sensitive and the most cold-tolerant ecotypes, suggesting the utility of this collection of natural variants in mining cold tolerance genes (Fig. 2d). To facilitate the identification of most of the cold tolerance genes, we (i) phenotyped the ecotypes for responses to cold stress multiple times; and we used data of each experiment as well as the mean from all experiments to conduct GWAS. Two models, Linear Regression (LR) and EMMAX, were assessed to accommodate population structures. We identified 33 candidate cold-responsive genes through GWAS (Table S4). Only 11 of the 33 genes were identified in GWAS when either LR or EMMAX model was used; 15 were identified in analyses with EMMAX and seven with the LR model (Table 1; Table S4). Analyses of at least two independent insertion mutants for 29 of these genes identified 16 cold tolerance genes (Table 1). Loss of function mutants of nine of the 16 cold tolerance genes with reduced transcript levels under cold stress showed enhanced cold tolerance; while, mutants of four genes induced during cold stress showed increased sensitivity to prolonged cold stress (Fig. 2e; Table 1). For three genes, an inverse relationship between the transcript levels and responses of mutants to cold stress was not observed. The inverse relationship between the growth of mutants and corresponding steady-state transcript levels of these 13 genes suggests that the majority of the 16 identified cold tolerance genes are regulated at the transcriptional level for adapting Arabidopsis to cold stress. In addition to the 16 cold tolerance genes, altered phenotypes for a single mutant of each of the nine genes were observed (Table 2). This suggests that these nine genes may have a subtle effect on cold tolerance. Investigation of additional mutants or complementation analyses of the loss-of-function mutants will establish the role of these nine putative cold tolerance genes.

Blast2GO analysis of all 16 cold tolerance genes revealed that the 16 genes could be grouped into 58 classes based on the biological processes they are involved in (Table 1; Fig. S4; Table S8-S10). This showed the complexity of cold tolerance mechanisms that interplay in adapting Arabidopsis to prolonged cold stress. We employed MapMan to map the identified 15 cold tolerance genes (Table1) and eight of the nine strong candidate cold-tolerance genes (Table 2) showing differential expression due to cold stress (Fig. S3; Table S5) onto metabolism, biotic stress, cellular response, proteasome, autophagy and cellular function categories (Fig. S8). MapMan analysis interestingly revealed the involvement of lipid metabolism (*ADS2*), biotic stress-related genes (*NB-ARC LRR, TIR-NB-LRR, AtRLP39, PER72, LRR protein kinase*), a protein involved in heat stress (DNAJ heat shock N-terminal domain-containing protein), ubiquitin and autophagy-dependent degradation, proteolysis (*EDA41*), vesicle transport and protein targeting (*AtSYP112*) and transcriptional regulation (*HMGB6*, stress-associated protein 7; *AtMYB42*) in response to cold stress (Fig. S8).

We identified a strong candidate cold-tolerance gene encoding a MYB transcription factor, *AtMYB42*, a homologue of *MYB15*, which was shown to be involved in cold tolerance earlier (Table 2). Overexpression of *MYB15* resulted in decreased freezing tolerance, while its knock-out mutant displayed an improved freezing tolerance (Agarwal et al., 2006). In our study, a T-DNA insertion *atmyb42* mutant showed enhanced cold tolerance (Table 2).

Transcription factors are involved in regulating the expression of the cold-responsive genes. For example, C-REPEAT BINDING FACTOR (CBF)-mediated cold signaling pathway (Chinnusamy et al., 2010; Jeon and Kim, 2013). Regulation of CBF genes plays a crucial role in the CBF-COR signaling pathway (Liu et al., 2019). The promoters of CBFs contain MYB recognition sequences suggesting MYB-related transcription factor participation in the cold induction of CBFs (Chinnusamy et al., 2003). MYB15 interacts physically with ICE1, which regulates the transcription of CBF genes in response to cold (Chinnusamy et al., 2003). Overexpression of ICE1 boosts the expression of the CBF regulon, thus improving freezing tolerance in transgenic plants (Chinnusamy et al., 2003).

Our findings in the present study (Fig. S8) also corroborate the earlier studies exhibiting the involvement of lipid metabolism, protein degradation, and the ubiquitin-proteasome system and DnaJ proteins in cold stress (Barrero-Sicilia et al., 2017; Kazemi-Shahandashti and Maali-Amiri, 2018; Xu and Xue, 2019). Lipid metabolism plays a key role in response to cold stress (Barrero-Sicilia et al., 2017). In cold stress, one of the adaptive responses is re-modeling cell membrane fluidity, which is achieved by increasing the unsaturated fatty acid composition of membrane lipids (Upchurch, 2008). Transcriptomics analysis of amino acid catabolism established a link between cellular regulation and protein degradation in response to a variety of environmental stresses, including cold stress (Kazemi-Shahandashti and Maali-Amiri, 2018). E3 ubiquitin ligases are involved with biotic and abiotic stresses including cold stress in Arabidopsis (Dong et al., 2006; Suh and Kim, 2015) and rice (Byun et al., 2017; Xu and Xue, 2019). The DnaJ proteins are treated as common cellular stress sensors because of their expression by many factors such as heat, cold, and drought (Rajan and D’Silva, 2009, Liu and He, 2020). The common pathways among cold stress and other abiotic and biotic stress signaling suggest the cross-talks among the pathways (Solanke and Sharma, 2008).

Identification of five novel genes encoding LRR domain containing proteins is worth noting. Temperature affects disease resistance by broadly influencing plant growth, regulating plant-pathogen interactions and defense responses mediated by several disease resistance (*R*) genes (Garrett et al., 2006). The role of NBS-LRR genes in freezing tolerance has been established (Huang et al., 2010; Yang et al., 2010; Zbierzak et al., 2013). NB-LRR receptor functions are known to be modulated by cold stress through the integration of an alternative H2A.Z histone variant into nucleosomes (Alcázar and Parker, 2011). NLRs or NLR-like proteins act as centers linking low-temperature stress and salicylic acid (SA)-dependent growth inhibition (Zbierzak et al., 2013). Over-expression of an LHY-CCA1-Like transcription factor SgRVE6 results in increased expression of 6 NB-LRR encoding genes associated with tobacco cold-tolerance, and it provides the transgenic tobaccos with higher tolerance to cold stress (Chen et al., 2020). Constant exposure to cold or low temperatures might result in the accumulation of SA and the suppression of development (Chen et al., 2015). Inactivation of a ubiquitin-conjugating enzyme, UBC13, compromises cold-responsive gene activation and causes elevated SA concentration and growth inhibition at low temperature. The phenotypes of the *ubc13* mutant are partially dependent on an NLR gene, *SNC1*, implying that UBC13 is engaged in NLR function during low-temperature stress (Wang et al., 2019). The defense regulator genes *SAG101, EDS1*, and *PAD4* negatively regulate the freezing tolerance in Arabidopsis, in part by modulating SA and diacylglycerol (DAG) homeostasis (Chen et al., 2015). The diacylglycerol acyltransferase 1 (DGAT1) is highly cold-responsive, SA downregulates its cold-responsive expression (Chen et al., 2015). DGAT1 catalyzes the final step in the triacylglycerol (TAG) assembly by acyl transfer from acyl-CoA to DAG. During cold acclimation, freezing tolerant plants displayed higher DGAT1 expression, which resulted in increased accumulation of TAG in response to subsequent freezing (Chen et al., 2015). DAG metabolism is also believed to act downstream of defense regulator genes SAG101, EDS1, and PAD4 in the SA-associated cold signaling pathway (Chen et al., 2015).

The chilling sensitive (*chs*) mutants, *chs2* and *chs3* of genes encoding R proteins of the TIR-NB-LRR class exhibited accumulation of high SA levels, specifically under cold stress (Eremina et al., 2089). None of the three NBS-LRR genes identified showed any similarity to previously cloned NB-LRR genes involved in freezing tolerance. *AT1G61310* encoding an NBS-LRR protein has been annotated to be a disease resistance-like protein. *AT5G41750* encoding a TIR-NB-LRR protein was previously shown to be a candidate for the *DM1* (*Dangerous Mix 1*) gene involved in autoimmunity and incompatibility response (Bomblies et al., 2007). It is becoming evident that NB-LRR proteins, LRR-kinase and RLP may have a significant role in signaling cold tolerance pathway in plants.

Several studies have indicated that abiotic stress signaling pathways overlap with the disease resistance signaling pathways (Lee and Yeom, 2015). Some of the NB-LRR-type *R* genes serve as non-immune receptors and involved in signaling for plant development. When grown below 16°C, the Arabidopsis *chilling-sensitive 2* (*chs2*) mutant demonstrated temperature-sensitive growth abnormalities comparable to those detected during defense responses (Huang et al., 2010). A gain-of-function mutant allele of the *RPP4* gene was detected in the *chs2* mutant. The mutant allele increases chilling sensitivity and expression of pathogenesis-related (*PR*) genes, the production of hydrogen peroxide and SA, when the mutant is cultured at 16°C (Huang et al., 2010). The Arabidopsis *chs3* mutant exhibiting induction of defense responses showed stunted growth and chlorosis at 16°C (Yang et al., 2010). *CHS3* encodes a TNL-LIM-type NB-LRR *R* gene regulates the freezing tolerance.

Assignment of cold tolerance function to 14 of the 16 identified genes is surprising. The overlap between the cold tolerance-related genes in prior and this study was observed just for two genes. One possible explanation could be that cold tolerance is regulated by a very complex process and identification of all components of this process is yet to be accomplished. We looked at the lesions that resulted in the mutant alleles for 16 genes involved in adaptation of the natural variants to the temperate climate of the northern hemisphere. For none of the genes, we observed a nonsense mutation. One possible reason for this is that the genes may have vital and multiple functions, as we have seen for many of the genes identified in this study. Natural selection shapes the expression levels or structure of the involved proteins or enzymes without compromising the other functions encoded by the genes during generation of new functions for adapting plants to new environments or growing conditions. We observed that SNPs identified by GWAS were localized to 5’-UTRs of three genes, 3’-UTR of one, and introns of three genes (Table 1; Table S4). A synonymous mutation was detected in *AT4G12000*, which encodes a SNARE-associated Golgi protein. This gene is highly expressed during cold stress, and two knockout mutants for this gene clearly showed super-sensitivity to cold stress (Table 1; Fig. S3m; Fig. S5f). It is possible that transcription of this gene could be impacted by the synonymous mutation if it is localized to a *cis*-acting element. Transcriptional regulation through *cis*-elements localized to UTRs is well established (Rose, 2019; Srivastava et al., 2018). Introns can contain splicing-regulatory sequences to autoregulate alternate splicing and transcription regulation (Thomas et al., 2012). Synonymous mutations in the open reading frames may also cause structural changes in mRNAs leading to changes in protein translation efficiency (Li et al., 2019a).

The GWAS, together with T-DNA insertion mutant analyses, revealed 14 novel cold tolerance genes and nine additional strong candidate cold tolerance genes. Only two of the 16 identified genes were previously identified. Nature shapes the expression of most of these genes without causing any changes in protein structures, presumably because of their multiple functions, some of which could be vital. Loss of function mutants for these genes presumably lack necessary fitness and are selected out by natural selection. The identified cold tolerance genes in this study provide a strong base for better understanding cold tolerance in plants and also developing crop varieties with resilience to cold stress, necessary for securing food supply under the unpredictable weather patterns resulting from climate change.

## MATERIALS AND METHODS

### Analyses of phenotypes

Seeds of 65 T-DNA insertion and one transposon-induced mutants for 33 candidate cold stress-related genes identified by GWAS were obtained from the Arabidopsis Biological Resource Center (https://abrc.osu.edu) and propagated directly under optimal greenhouse conditions to obtain sufficient seeds for this study. Knockout mutants were characterized for homozygosity as described in the following sections. Leaf samples were collected for genomic DNA isolation from each of the ten individual plants of a knockout mutant. At least two knockout mutants were characterized for homozygosity and studied for each candidate gene (Fig. 1e-f; Fig. S4; Table S6-S7). Knockout mutants and wild-type Col-0 ecotype were phenotyped under control and prolonged cold-stress conditions as described in the Results section to determine if any of the 33 putative cold stress-related genes play a role in cold tolerance. Two-dimensional images of the rosette leaves of individual mutants were captured and analyzed by Matlab GUI software (Method S2). The digital images were converted to pixel data for determining the relative rosette leaf growth rate of the mutant compared to wild-type Col-0 under control (C) and cold stress (CS) conditions.

### Genome-wide association studies

For GWAS, the average trait (changes in the rosette area in extended cold stress) value of each accession’s biological replicates was taken. The GWA analysis was performed in the easyGWAS web interface (Grimm et al., 2017) using the linear regression (LR) model or Efficient Mixed-Model Association eXpedited (EMMAX).

### Validation of putative genes by reverse genetics approach

To validate the cold-tolerance function of the putative genes identified by GWAS, we studied at least two independent T-DNA insertion mutants for each of the candidate genes. The information about primer sequences, insertion locations, and the estimated T-DNA confirmation product size was obtained from the T-DNA Primer Design site (http://signal.salk.edu/tdnaprimers.2.html) (Table S6). The homozygous plants for any T-DNA insert from individual segregants were identified essentially by a two-step PCR genotyping assay (O’Malley et al., 2015).

### Screening of T-DNA Insertion Lines

A gene/genome-specific primer (GSP: LP, RP - Left, right genomic primer) pair spanning the predicted T-DNA insertion site was used for the first PCR reaction to detect the presence of a wild-type copy (WT copy) of the gene in the wild type or heterozygous individuals. However, no band was amplified for a homozygous plant because both copies of the gene contain the T-DNA insert, whose large size precludes PCR amplification. The lack of a wild-type gene-specific PCR provided strong evidence that the line is homozygous for the insert (Fig. 1e). The second PCR reaction was used to validate that the homozygosity for a T-DNA insert in the gene. In Col-0, we failed to amplify the T-DNA inserted genomic region in the second PCR, while a T-DNA and target insertion site-specific PCR product was amplified in the heterozygous and homozygous T-DNA insertion mutants (Fig. 1e). The homozygous lines showed lack of the gene-specific and presence of T-DNA insertion site-specific PCR amplification. The heterozygotes showed amplification of both types of PCR products; i.e., gene-specific and T-DNA and insertion site-specific (Fig. 1e). The T-DNA insertion site was selectively amplified, using a combination of a left border primer (LB - the left T-DNA border primer) and the correctly oriented GSP primer (LP or RP) specific to the target insertion site (Fig. 1e).

### BLAST search analysis

Blast2GO was used to determine the function and localization of the candidate genes. Blast2GO is a widely used annotation platform that uses homology searches to associate sequence with Gene Ontology (GO) terms and other functional annotations (Conesa et al., 2005; Götz et a. 2008). Blast2GO generated Gene Ontology annotations for the three sub-trees of GO, (a) Biological process, (b) Molecular Function, and (c) Cellular Component.

### MapMan Analysis

With an objective to display cold stress-responsive genes onto pathways, the MapMan (Usadel et al., 2009) was used to analyze the 16 cold tolerance genes (Table 1) and nine strong cold tolerance genes (Table 2) that are induced or suppressed following cold stress (Fig. S3; Table S5).

### Statistical analysis

Using package R program version 1.6.1 (Ihaka and Gentleman, 1996), the Student’s t-test was performed to determine the significant difference of comparative growth rate of mutants to Col-0 between control and prolonged cold-stress conditions while one-way ANOVA (Analysis Of Variance) was performed for determining growth rate differences between the ecotypes.

## SUPPLEMENTAL INFORMATION

**Figure S1. (a)** Geographical locations of 417 Arabidopsis ecotypes used in this study.

**(b)** QQ Plot of observed versus expected *p*-values for the changes of leaf area under prolonged cold stress GWAS analysis for all SNPs.

**(c)** Frequency distribution of the 417 accessions for proportionate cold tolerance. The location of reference accession Col-0 is indicated with a red arrow. Growth rate data for each accession are given in Table S2. The growth rate of each ecotype (%) is calculated as growth at termination of exposure to cold stress (on 30th day of treatment) X 100/Growth before initiation of treatment (0th day of treatment). Proportionate tolerance of each ecotype is calculated as the growth rate of each ecotype X 100/ the summation of growth rates of 417 ecotypes (detailed information is on Supplemental Fig. S2).

**Figure S2.** Differential tolerance of 417 Arabidopsis ecotypes to continuous cold stress.

**Figure S3.** Expression patterns of sixteen identified cold-responsive genes **(a-p)** during exposure to cold stress using the eFP database (Winter et al., 2007; http://bar.utoronto.ca/efp2/Arabidopsis/Arabidopsis_eFPBrowser2.html).

**Figure S4.** Differential tolerance of T-DNA mutants of the identified genes to continuous cold stress. Here, a few representative cold-tolerant (PYL-6, Gr-1, and Kil-0) and cold-sensitive (Zdr-1) ecotypes are shown.

**Figure S5.** Nine regions of interest containing genes that contribute towards cold tolerance. **(a–i)** Each panel shows data for a genomic region of interest for which the mutant analysis uncovered cold stress-responsive genes. On the left, output plot of *p*-values (-log base 10) in a 5-kb window for association of SNPs with phenotypic variation, obtained from easyGWAS is presented. On the right, rosette leaf growth rates of mutants with respect to Col-0 are presented. The relative rosette leaf growth rate in the mutant compared to wild-type Col-0 is significantly different in control (C) and cold stress (CS) (*p* < Bonferroni adjusted α) conditions. C, Comparative growth rate of the mutant relative to wild-type Col-0 in control condition; CS, Comparative growth rate of the mutant relative to wild-type Col-0 in cold stress.

**Figure S6.** Gene Ontology (GO) analysis. The pie graphs showing the grouping of 16 cold-response genes to (a) 58 classes based on biological processes, (b) 17 classes based on molecular functions, and (c) 13 classes based on their subcellular locations or as cellular components. The numbers in parentheses show the percentage of total genes in each functional categorization of genes. The summary of genes in each functional categorization is represented in Table S8-S10.

**Figure S7.** The KEGG pathways showing involvement of *AT2G18260* (EC:2.7.7.3) in pantothenate and CoA biosynthesis pathway (a), *AT2G31360* (EC:1.14.19.1) in the biosynthesis of unsaturated fatty acids (b), *AT2G27120* (EC:2.7.7.7) in DNA replication, base/nucleotide excision repair, and purine metabolism pathways (c-f), *AT3G53520* (EC:4.1.1.35) involved in amino sugar and nucleotide sugar metabolism pathways (g) and *AT5G54960* (EC:4.1.1.1) involved in glycolysis/gluconeogenesis (h).

**Figure S8:** Overview of differentially regulated genes involved in different metabolic processes. Gene transcripts that are induced or repressed as a result of cold stress are shown for (a) metabolism overview, (b) biotic stress, (c) cellular response overview, (d) ubiquitin and autophagy-dependent degradation, (e) cell functions overview, (f) large enzyme families overview, (g) cell wall precursors, (h) Glycolysis-TCA, (i) RNA-protein synthesis. The figure was generated using the MapMan visualization tool (Usadel et al., 2009) on genes for which differential expression values were available (Table-1 & 2).

**Method-S1.** Correctly register image.m

**Method-S2.** Matlab GUI

**Table-S1.** Details of 417 Arabidopsis ecotypes used for phenotyping and analysis

**Table-S2.** The growth rate of 417 Arabidopsis ecotypes under cold stress.

**Table-S3.** Proportionate tolerance* of 417 Arabidopsis ecotypes under prolonged cold stress.

**Table-S4.** SNPs associated with responses to cold stress in Arabidopsis

**Table S5.** Expression patterns of 32 putative cold-responsive genes during exposure to cold stress using the eFP database (Winter et al., 2007; http://bar.utoronto.ca/efp2/Arabidopsis/Arabidopsis_eFPBrowser2.html).

**Table-S6.** Primers used for analyzing the T-DNA insertion lines of 33 candidate cold-stress responsive genes.

**Table S7.** Phenotypes of T-DNA insertion mutants for each of the 33 candidate cold-stress responsive genes identified in GWAS (Table S1).

**Table S8.** Gene Ontology (GO) annotations for Biological process.

## ACKNOWLEDGMENTS

We thank David Grant for reviewing the manuscript and his comments. This work was supported by a Plant Sciences Institute, Iowa State University grant to MKB. We are also thankful to Yin Lab members and Trevor Nolan from the Department of Genetics, Development and Cell Biology, Iowa State University, for helping in increasing seeds for the experimentation.

## COMPETING INTEREST

The authors have declared that no competing interests exist.

